# Ultrasonic Reporter of Kinase Activity

**DOI:** 10.64898/2025.11.30.691048

**Authors:** Jee Won Yang, Zhiyang Jin, Ting-Yu Wang, Mikhail G. Shapiro

**Author notes:** Corresponding authors: MGS.

## Abstract

Protein kinases are an essential class of enzymes that regulate cellular signaling pathways, with their dysregulation implicated in pathologies such as cancer and neurodegenerative diseases. Despite the existence of high-performance fluorescent biosensors of kinase activity, it remains challenging to study the function and regulation pathways of kinases in opaque tissues due to the limited tissue penetration of light. To address this limitation, we introduce the first ultrasonic reporter of kinase activity (UReKA), designed to detect protein kinase A (PKA) activity by altering the ultrasound scattering of gas vesicles, a unique class of air-filled protein nanostructures. We engineer a gas vesicle shell protein to respond to PKA, demonstrate the functionality of UReKA both in purified protein format and expressed in mammalian cells, and showcase its capacity to monitor PKA signaling in response to pharmacological stimulation or genetic mutation. This work makes it possible to visualize cellular functional activity in opaque media, with broad potential for future applications in cancer biology, cellular development, and drug discovery.

## INTRODUCTION

Kinases are a diverse family of enzymes that regulate a wide range of biological processes across virtually all cell types through phosphorylation^1–3^, including signal transduction^4^, metabolic homeostasis^5^, immune regulation^5^, and neuroprotection^6^. Moreover, aberrant kinase signaling is implicated in numerous pathologies such as cancer, chronic inflammation, diabetes, and neurological disorders^6–8^, making this class of enzymes prime targets for therapeutic intervention^8,9^. Hence, the ability to visualize kinase activity with high spatiotemporal precision is increasingly needed to study the dynamics of cellular signal transduction and facilitate drug development and therapeutic monitoring.

Due to the central role of kinases in cellular function and dynamics, many imaging and analysis methods have been developed to understand their activity, substrate specificity, and function. However, traditional protein kinase assays including phosphoproteomics^10^ and radiolabeled [P32] techniques^11^ require cell lysis, preventing real-time observation of dynamic phosphorylation events. More recently, genetically encoded biosensors based on fluorescence, luminescence, Förster resonance energy transfer^12^, and fluorescence lifetime imaging microscopy techniques have greatly improved spatiotemporal resolution in live cells^13,14^ and animal models^15–17^. Yet, these optical techniques are inherently restricted by light scattering in tissues^18^, limiting their applicability to study kinase activity in opaque samples. While the advent of a noninvasive kinase reporter based on magnetic resonance imaging^19,20^ provided an alternative modality with greater tissue penetration, the system was not expressed in cells or demonstrate intracellular enzyme activity sensing.

In contrast, ultrasound offers unique advantages for imaging cellular responses due to its ability to penetrate centimeters into opaque tissue while maintaining high spatial (∼100 µm) and temporal (∼1 ms) resolution^21,22^. Genetically encoded ultrasound contrast agents have been developed from gas vesicles (GVs), air-filled protein nanostructures derived from buoyant microbes^23^. These agents function as acoustic reporter genes^24–26^ and biosensors of protease activity^27^ and intracellular calcium^28^, with their acoustic properties regulated by GvpC, an α-helical surface protein that stiffens the GV shell and controls buckling, collapse pressure, and nonlinear contrast generation^29–33^. In this study, we introduce UReKA, the first acoustic biosensor for kinase activity, engineered to enable dynamic, noninvasive monitoring of intracellular PKA activity.

## RESULTS

### UReKA Design and Identification

To demonstrate the feasibility of UReKA, we selected PKA as a model sensing target due to its involvement in diverse cellular pathways^34–37^. We hypothesized that phosphorylation-induced charge and conformational changes within a surface-binding protein could modulate the mechanical properties of GVs, leading to a change in nonlinear ultrasound contrast (**Fig 1a**). To test this concept, we engineered variants of GvpC, a structural protein known to mechanically reinforce the GV shell, to incorporate various PKA substrate motifs^38,13^. We hypothesized that phosphorylation of the embedded serine/threonine residues would reduce GvpC’s affinity for the GV surface, resulting in a more deformable shell and a corresponding enhancement in nonlinear acoustic response^30^.

**Figure 1.**
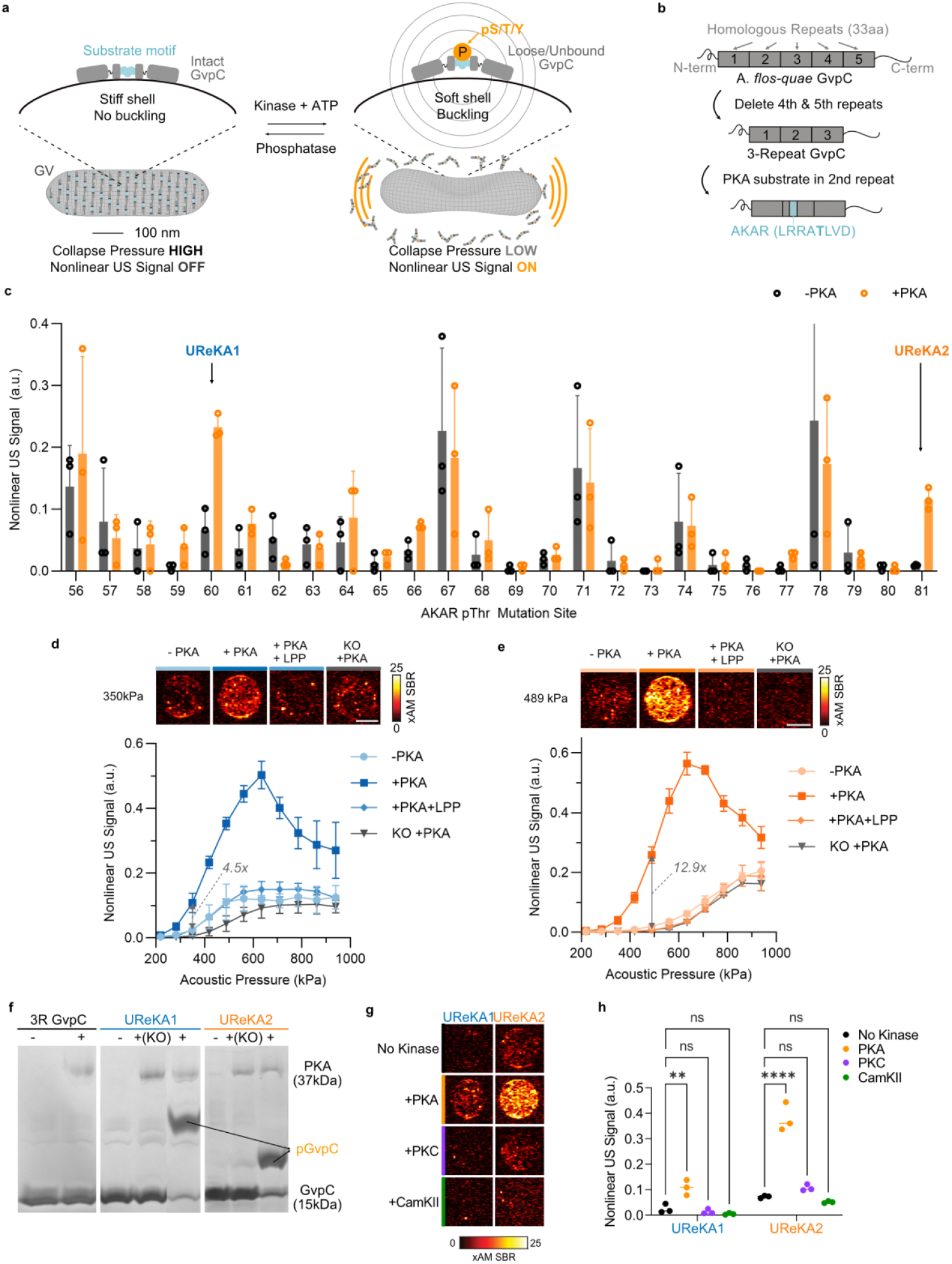
Identification and Characterization of UReKA. a) General design schematics of UReKA and its proposed mechanism of action. B) Schematics of UReKA variant GvpC design process. c) Nonlinear ultrasound normalized by B-mode signal screening of AKAR-UReKA variants at various pThr position (56-81) in 3R GvpC at 419 kPa. Significance determined by multiple t-test (n=3). d) (Top) Representative ultrasound images of agarose phantom containing UReKA1 and nonlinear ultrasound signal at 350 kPa (4V). Scale bar=1mm. (Bottom) Nonlinear ultrasound signal normalized by B-mode intensity as a function of acoustic pressure for UReKA1, without PKA, with PKA, with PKA and LPP, with KO control (n=3). e) (Top) Representative nonlinear (xAM) ultrasound images of agarose phantom containing UReKA2 and nonlinear ultrasound signal at 489 kPa. Scale bar=1mm. (Bottom) Nonlinear ultrasound signal normalized by B-mode intensity as a function of acoustic pressure for UReKA2, without PKA, with PKA, with PKA and LPP, with KO control (n=3). f) Coomassie Brilliant Blue-stained SuperSep™ Phos-Tag® SDS-Page gel image of UReKA1 and UReKA2 with respective KO controls incubated with and without PKA. g) Representative ultrasound images of agarose phantom containing UReKA1 and UReKA2 at 350 kPa and 489 kPa, respectively, incubated with PKA, PKC, CamKII. Scale bar=1mm. h) Nonlinear ultrasound signal normalized by B-mode intensity as a function of acoustic pressure for UReKA1 and UReKA2 at 350 kPa and 489 kPa, respectively, with PKA, PKC, CamKII. Scale bar=1mm. (n=3)

Conversely, we envisioned the activity of a phosphatase reversing phosphorylation, returning GvpC to the GV shell, and enabling reversible imaging of phosphorylation dynamics.

Based on previous experience in GvpC engineering^26,27^, we employed a three-repeat GvpC (3R-GvpC) construct as the backbone, in which the last two of the five 33-amino acid homologous regions in the wild-type Ana GvpC gene were deleted (**Fig 1b**). A library of GvpC variants was generated by inserting the AKAR PKA substrate sequence (LRRATLVD) ^13^ into individual positions within the second repeat (residues 56– 81), replacing the wild-type amino acids at the corresponding positions. Screening of these constructs revealed position-dependent changes in nonlinear ultrasound intensity following incubation with and without PKA (**Fig. 1c**). Among the variants, two designs—UReKA1 (with the cognate threonine at position 60 - Thr60) and UReKA2 (Thr81)—exhibited the largest phosphorylation-dependent signal enhancement under cross-propagating amplitude modulation (xAM) ^30^ nonlinear imaging at 419 kPa peak positive transmit pressure, which were then selected for detailed characterization.

Both UReKA1 and UReKA2 displayed reversible, phosphorylation-dependent modulation of nonlinear ultrasound contrast. UReKA1 showed a 4.5-fold signal increase upon PKA treatment when imaged at 350 kPa, fully reversed by lambda-protein phosphatase (LPP), while UReKA2 demonstrated a 12.9-fold increase when imaged at 489 kPa (**Fig. 1d–e**). In both cases, point mutation of the threonine to alanine (KO mutant) abolished the response, confirming that the observed modulation arose from specific phosphorylation. These acoustic measurements correlated with pressurized absorbance spectroscopy assays^39^, where the midpoint collapse pressure decreased from 398 ± 43 kPa to 227 ± 20 kPa for UReKA1 and from 424 ± 64 kPa to 283 ± 15 kPa for UReKA2 upon PKA phosphorylation (**Supplementary Fig. 1a–c**). The LPP co-incubation restored collapse pressure to baseline, supporting the mechanical reversibility of the system. Moreover, Phos-tag SDS-PAGE^42^ further verified phosphorylation of GvpC upon PKA treatment (Fig. 1f).

To verify the sensors’ kinase specificity, we tested both UReKA variants against PKC and CaMKII, two serine/threonine kinases with relatively similar consensus sequences to PKA^43^. Neither enzyme induced detectable changes in collapse pressure or nonlinear contrast (**Figs. 1g–h, Supplementary Figs. 1d–e**), confirming selective responsiveness to PKA.

After identifying AKAR-based UReKA as a robust, reversible reporter, we explored strategies to enhance sensitivity and signal dynamics. Insertion of the AKAR motif at alternative repeat positions, or in multiple repeats, disrupted GvpC–GV binding and failed to produce functional constructs (Supplementary Fig. 2a). Similarly, fusion of the phosphoamino acid–binding FHA1 domain^40,41^ to AKAR-GvpC was hypothesized to amplify conformational change but instead led to protein aggregation and loss of function (**Supplementary Fig. 2b**). We also screened Kemptide (LRRASLG) and modified Kemptide (LRRASLP) substrates (**Supplementary Fig. 2c-d**) and demonstrating that the design principles can be generalizable with alternative substrate sequences, despite the reduced dynamic range and weaker GV reinforcement compared to AKAR-based variants. Collectively, these results establish UReKA1 and UReKA2 as our best constructs for the first kinase-responsive acoustic biosensors, exhibiting strong, reversible, and selective ultrasound readouts of enzymatic phosphorylation.

### Characterization of UReKA in Purified GVs

To understand the molecular basis of UReKA activation, we first verified the specific phosphorylation site introduced into the GvpC. Mass spectrometry confirmed a single phosphorylated residue at Thr60 within the AKAR motif of UReKA1, with no other phosphorylation detected across the protein sequence (**Fig. 2a**). These data validated that PKA specifically modifies the engineered site without off-target modification of native serine or threonine residues in GvpC.

**Fig 2:**
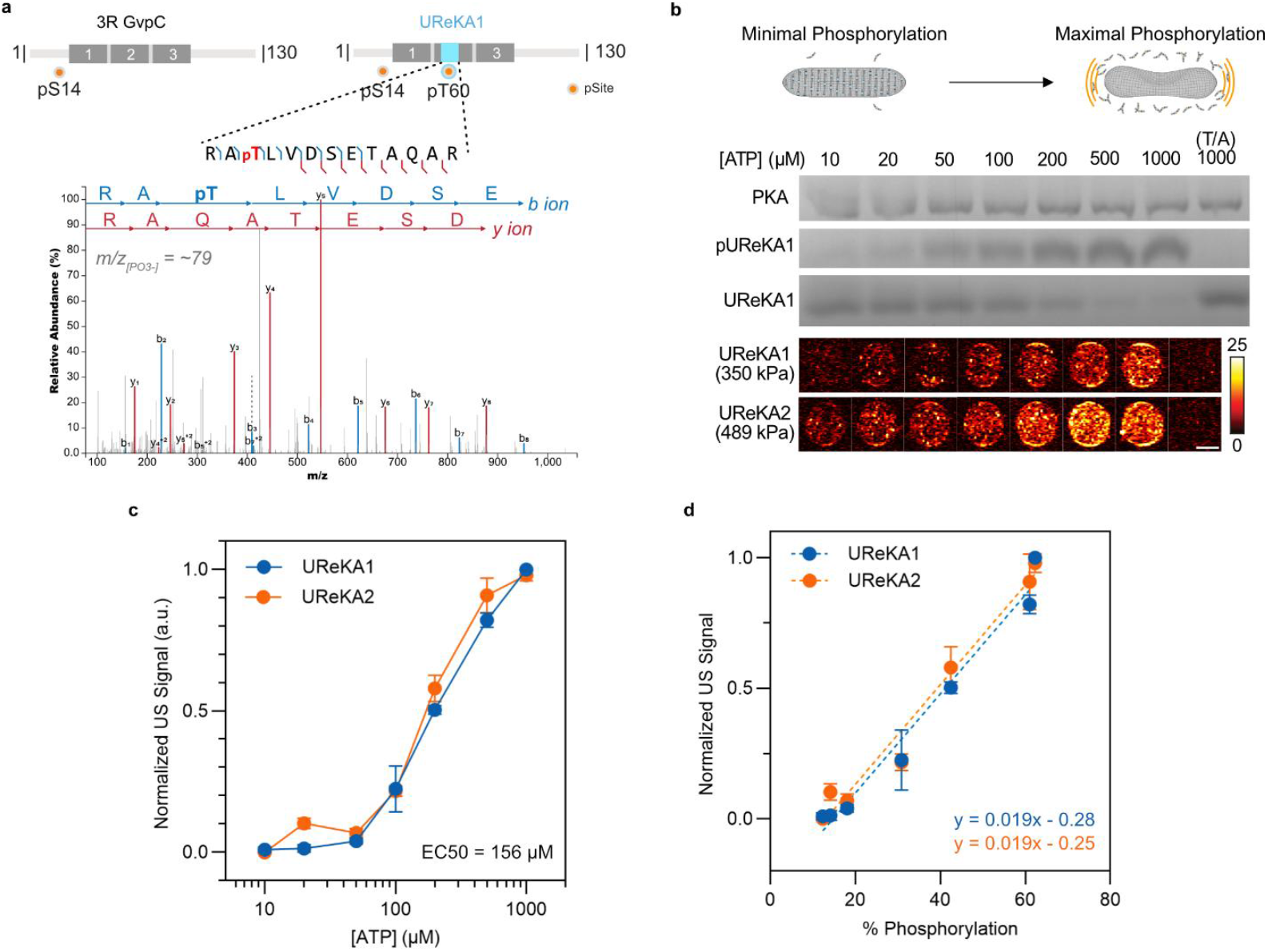
Quantitative Analysis of UReKA Phopsphorylation. a) Mass spectrometry schematics. Annotated MS/MS spectrum of the phosphorylated peptide in UReKA1. Matched b- and y-ions are color-coded in blue and red, respectively. The primary amino acid sequence of the identified phosphopeptide is shown, with all matched fragment ions labeled. The p preceding residues denotes phosphorylation. b) (Top) Coomassie Brilliant Blue-stained SuperSep™ Phos-Tag® SDS-Page gel image with various concentration of ATP. (Bottom) Respective xAM images of UReKA1 (at 350 kPa) and UReKA2 (at 489 kPa). c) Normalized nonlinear signal of UReKA1 (blue) and UReKA2 (orange) as a function of ATP concentration. n=3 biological replicates. The legend lists the EC50 of UReKA determined from fitting a 4-parameter logistical (4PL) model. d) Normalized ultrasound signal of UReKA1 (blue) and UReKA2 (orange) as a function of percent phosphorylated fraction of UReKA determined by mass spectrometry. n=3 biological replicates.

We next quantified the relationship between the degree of phosphorylation and acoustic response by titrating ATP concentrations during PKA incubation (**Fig. 2b**). As ATP concentration increased from 10 µM to 1 mM, Phos-Tag SDS-PAGE revealed a progressive upward shift, corresponding to incremental phosphorylation of UReKA1 and UReKA2. The nonlinear ultrasound contrast rose proportionally with increasing ATP, reaching saturation at 500–1000 µM. Fitting a four-parameter logistic model yielded an EC_50_ of 156 µM, well below physiological intracellular ATP levels (∼3 mM) ^45^, indicating that UReKA can operate within biologically relevant regimes (**Fig. 2c**). As expected, parallel xAM ultrasound signal curves obtained across all ATP concentrations showed a monotonic increase in nonlinear signal amplitude with phosphorylation (**Supplementary Fig. 3a–b**).

Moreoverquantitative mass spectrometry analysis determined that UReKA1 and UReKA2 exhibited a linear correlation (R^2^ > 0.97) between phosphorylation extent and acoustic signal intensity (**Fig. 2d**), where the half-maximal ultrasound signal corresponded to 40 ± 5 % phosphorylation of GvpC. This linear scaling indicates that the nonlinear ultrasound response directly reflects the fraction of phosphorylated GvpC bound to the gas vesicle surface, thereby allowing quantitative readout of kinase activity *in vitro*. These data establish that UReKA’s acoustic contrast varies continuously with phosphorylation state, providing a tunable and quantitative correlation between enzymatic modification and mechanical modulation of nanoscale acoustic reporters.

### Characterizing Molecular Mechanism and Kinetics

Having established the specificity and reversibility of UReKA’s phosphorylation response, we next investigated the kinetic behavior and underlying molecular mechanism governing its activation. Time-course experiments revealed that the nonlinear ultrasound signal of UReKA1 and UReKA2 increased gradually following PKA incubation, reaching maximal contrast after approximately 8 h and 12 h, respectively (**Fig. 3a–b**). The acoustic signal first became detectable after 2–3 h, whereas Phos-Tag SDS-PAGE analysis showed that phosphorylation of the GvpC scaffold was already complete within this early period (**Fig. 3c, Supplementary Fig. 4a**). The decoupling between phosphorylation and acoustic activation thus suggests that the rate-limiting step in UReKA response is not the enzymatic modification itself, but rather a slower structural transition or dissociation process that follows phosphorylation.

**Fig 3.**
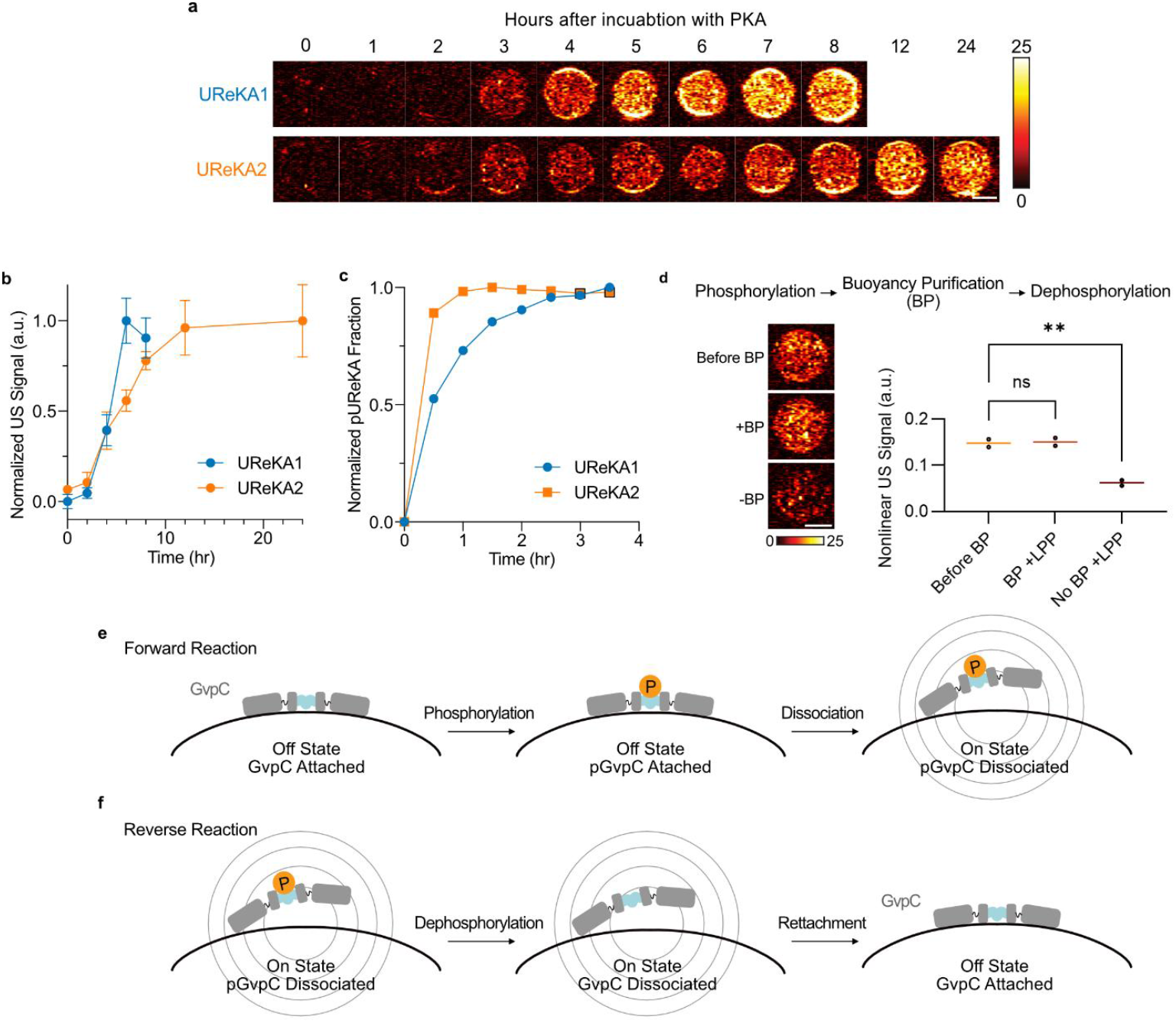
Kinetics & Molecular Mechanism of UReKA. a) xAM images of UReKA1 and UReKA2 incubated with PKA at various timepoints at 634kPa (for maximal signal). b) Normalized nonlinear US signal of UReKA1 (blue) and UReKA2 (orange) over time (n=3). c) Normalized phosphorylated fraction of UReKA1 (blue) and UReKA2 (orange) over time (n=1). d) Nonlinear ultrasound images (left) and signal (right) of UReKA1 incubated with PKA and phosphorylated UReKA1 incubated with LPP with or without buoyancy purification. (n=2). e-f) Schematics of the proposed molecular mechanism of UReKA with reversible phosphorylation reaction for the phosphorylation as forward reaction (e) and dephosphorylation of GvpC as reverse reaction (f).

To distinguish whether the signal enhancement arises from partial structural rearrangement or full detachment of phosphorylated GvpC from the gas-vesicle surface, we performed a buoyancy purification (BP) assay to separate the buoyant GV-bound fraction from the unbound GvpC (**Fig. 3d**). When phosphorylated UReKA1 samples were dephosphorylated by LPP before BP, both nonlinear ultrasound signal and collapse pressure were fully restored to baseline, consistent with previously shown data. In contrast, when LPP was applied after BP—acting only on the GV-bound fraction—the ultrasound signal remained elevated and indistinguishable from phosphorylated controls, indicating that the phosphorylated GvpC had already dissociated from the GV shell. Together, these data support a model in which phosphorylation induces GvpC detachment rather than an internal conformational shift within the bound state.

From these observations, we propose a two-step molecular mechanism for UReKA activation (**Fig. 3e–f**). In the forward reaction, phosphorylation of GvpC by PKA introduces negative charge that weakens electrostatic interactions with the GV surface, promoting dissociation and generating the acoustically “on” state. In the reverse reaction, dephosphorylation by LPP allows unmodified GvpC to re-associate with the GV shell, restoring mechanical rigidity and the acoustically “off” state. The observed delay between phosphorylation and acoustic activation therefore reflects the kinetics of protein–shell dissociation and reattachment, rather than chemical turnover at the phosphorylation site.

Altogether, these results establish that UReKA functions through a phosphorylation-triggered GvpC detachment mechanism, where enzyme-controlled modulation of GvpC-shell binding governs the mechanical and acoustic state of the reporter. This mechanistic understanding provides a foundation for further tuning of sensor response speed and reversibility through protein-interface engineering.

### In cellulo response of UReKA

Having validated UReKA’s reversible and quantitative response to phosphorylation *in vitro*, we next sought to demonstrate its functionality as an intracellular kinase activity sensor in mammalian cells. We selected UReKA1 as the prototype variant for *in cellulo* expression owing to its faster activation kinetics compared to UReKA2. To enable heterologous gas vesicle (GV) formation and UReKA expression, we employed a two-plasmid system in HEK293T cells (**Fig. 4a**). The first plasmid encoded the GV structural operon (*gvpNJKFGWV*) under a CMV promoter, while the second plasmid carried *gvpA* and the UReKA-modified *gvpC*, linked by an internal ribosome entry site (IRES) ^49^ element to control relative expression levels^25^. This modular configuration allowed fine-tuning of the GvpA:GvpC stoichiometry, which we hypothesized to be a critical determinant of UReKA activation dynamics due to the rate-limiting GvpC dissociation step.

**Fig 4.**
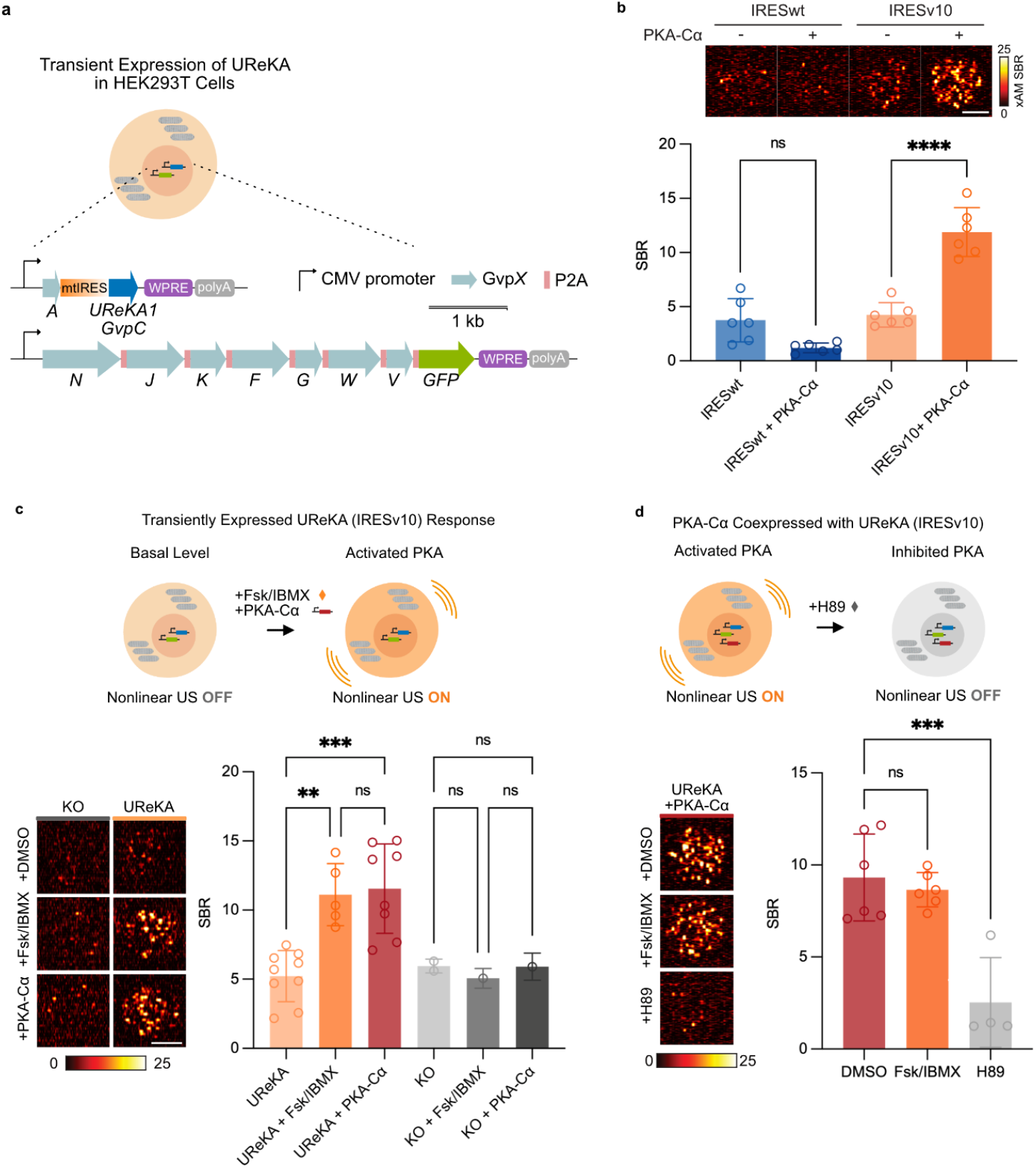
Pharmacological response of UReKA-GVs in HEK293T. a) Schematics of the genetic constructs for transient expression of UReKA in HEK293T cells and xAM images of transiently expressed UReKA in HEK293T cells with various versions of IRES mutant for decreased expression of UReKA-GvpC. IRESv10 mutant (UReKA-Iv10, blue star) was selected for all following expressions of UReKA in mammalian cells. b) (Top) Representative nonlinear xAM image at 419 kPa (top) of UReKA GVs with IRESwt and IRESv10 upstream of GvpC with or without co-expression of PKA-Cα catalytic subunit with respective SBR (below). n=6. c) (Top) Schematics of applying UReKA-Iv10 to image intracellular PKA activation through either forskolin/IBMX (Fsk/IBMX) stimulation or coexpression of PKA-Cα catalytic subunit. (Bottom) xAM CNR (left) and representative image (right) of UReKA-Iv10 with Fsk/IBMX stimulation or PKA-Cα coexpression at 419 kPa. T/A mutant was used as the KO control and DMSO was used as vehicle control. d) (Top) Schematics of transiently expressed UReKA-Iv10 coexpressed with PKA-Cα applied to image intracellular PKA inhibition with H89. (Bottom) xAM SBR (left) and representative image (right) of UReKA-Iv10 with Fsk/IBMX or H89 stimulation at 419 kPa. DMSO was used as vehicle control.

To optimize the GV–GvpC balance, we screened IRES mutants of varying translation efficiencies^50^ (**Supplementary Fig. 5**). Among these, the IRESv10 variant provided an optimal dynamic range—producing sufficient GvpC for structural reinforcement while avoiding excess unbound protein that could potentially buffer phosphorylation effects. Constructs using IRESv10 displayed markedly higher nonlinear ultrasound signal upon PKA activation compared to wild-type IRES, while maintaining minimal basal background (**Fig. 4b**). Consequently, the UReKA1-Iv10 variant was selected for all subsequent mammalian experiments.

We next evaluated UReKA’s response to intracellular modulation of PKA activity. Cells expressing UReKA1-Iv10 were stimulated with a combination of forskolin (FSK) ^51–53^, an adenylate cyclase activator, and IBMX^53^, a phosphodiesterase inhibitor, both of which elevate intracellular cAMP and thereby activate PKA (**Fig. 4c**). Alternatively, to mimic pathological PKA hyperactivation^55^, cells were co-transfected with a *PRKACA* construct encoding a constitutively active PKA catalytic subunit (PKA-Cα_L206R_). In both conditions, UReKA-expressing cells exhibited robust increases in nonlinear ultrasound contrast of 5 to 10 dB higher signal-to-background ratio (SBR) compared with vehicle-treated or kinase-inactive controls—demonstrating successful intracellular phosphorylation detection.

Conversely, treatment with the PKA inhibitor H89^54^ effectively reversed the signal enhancement (**Fig. 4d**), reducing the nonlinear response to baseline levels in PKA-Cα– overexpressing cells. Together, these results confirm that UReKA accurately reports changes in intracellular PKA activity with high sensitivity and reversibility. Overall, this system establishes UReKA as a genetically encodable, enzyme-responsive acoustic reporter capable of resolving kinase activity states within living mammalian cells. The tunable IRES-based design further provides a generalizable strategy for balancing stoichiometric expression of structural and functional protein components in acoustogenetic constructs.

## DISCUSSION

In this study, we introduced and validated UReKA as the first acoustic biosensors of kinase activity. Using purified recombinant GVs, we demonstrated that UReKA specifically responds to phosphorylation at engineered sites by the targeted kinase, with these modifications reversibly changing the the GVs’ nonlinear acoustic contrast. We also confirmed that the observed nonlinear signal activation is driven by detachment of GvpC from the surface of the GVs, similar to URoCs^27^. Additionally, we showed that UReKA expressed in mammalian cells effectively detects intracellular PKA activity, exhibiting increased or decreased signal in response to kinase activation or inhibition through pharmacological stimulation or genetic modification. Taken together, these results lay the groundwork for the first prototype of an ultrasonic reporter system for kinases that is quantitative and reversible.

The *in cellulo* applications demonstrated in this work suggest that UReKA has strong potential for future use in drug screening, especially for kinase inhibitors, and kinase activity imaging in a living animal for cancer and neurobiological research. The potential ability to monitor real-time kinase dynamics and pharmacological responses to novel therapeutics could provide valuable insights into cellular signaling pathways and drug mechanisms of action. Furthermore, complementary to fluorescence- and PET-based reporters, UReKA’s acoustic signal readout could in the future enable deep-tissue imaging and real-time molecular monitoring, offering a unique advantage for studying dynamic cellular signaling in its native physiological context.

While the genetic constructs described in this study could be applied to some experimental settings, further optimization is required to enhance UReKA’s adaptability and suitability for *in vivo* use as a live monitoring sensor. One key area for improvement is accelerating the sensor’s response kinetics beyond the hour-scale observed in this study. Following the examples from generations AKARs that improved over the past couple decades, future work could optimize UReKAs through various approaches including rational design, directed evolution, and computational modeling of specific substrate sequences^58,59^. For example, engineering a sensor that does not rely on complete GvpC dissociation may enable a faster turnover rate between different phosphorylation state and improve real-time monitoring of kinase activity. Additionally, although this study demonstrates robust signal dynamics in cellulo, further validation in tissue models and live animal system in a preclinically relevant scenario is needed to assess performance under complex physiological conditions. Another important direction is to develop UReKA variants for other target kinases such as PKC, Akt, or RTKs with enhanced sensitivity, which would enable broader and more reliable signal detection for a more diverse applications in biology and medicine.

Overall, UReKA represents the first approach to ultrasound-based molecular imaging for post-translational protein modification, providing a non-invasive, quantitative, and reversible method for monitoring kinase activity in real time.

This advance offers a ‘Eureka’ moment for molecular imaging of live-cell signals in opaque media.

## MATERIALS AND METHODS

### Design and cloning of genetic constructs

All plasmids used in this study were constructed using a combination of polymerase chain reaction (PCR) with Q5 polymerase and either Gibson Assembly^60^ or KLD mutagenesis, using enzymes from New England Biolabs (NEB) and custom primers from Integrated DNA Technologies (IDT). For screening and characterization of purified proteins, all *gvpC* gene sequences were codon-optimized for *E. coli* expression and introduced by substitution-insertion into the second repeat of the WT *gvpC* gene—from *Anabaena flos-aquae*—sequence in a pET28a expression vector (Novagen) driven by a T7 promoter and lac operator. Specifically, the construct encoding the three-repeat *Ana gvpC* gene was generated by deleting the fourth and fifth repeats from WT *Ana gvpC* sequence while appending a C-terminal 6xHis-tag (Addgene #85732) via KLD mutagenesis. Subsequently, codon-optimized PKA recognition motifs, including AKAR (LRRATLVD), kemptide (LRRASLG), or modified kemptide (LRRASLP) sequences, were introduced through KLD mutagenesis to generate the UReKA GvpC variants. Additional modifications, such as FHA1-fusion or SUMOylation-tag insertion, were achieved through Gibson Assembly.

For transient expression of UReKA in mammalian cells, all sequences were codon-optimized for human cell expression and cloned into pCMV-Sport vector with WPRE-hGH polyA driven by a CMV promoter. Similar modifications were made to the WT *gvpC* sequence generate to UReKA1 and control gvpC constructs, which were then inserted into an existing plasmid encoding the *gvpA* gene (Addgene #197588), preceded by an IRES sequence, using Gibson Assembly. An existing plasmid was used for the expression of all seven other chaperone proteins (*gvpNJKFGWV*) required for gas vesicle formation (Addgene #197589). For constitutive PKA activation, a gBlock encoding the *PRKACα* gene (NM_002730.4) was custom ordered from IDT and inserted into the backbone plasmid via Gibson Assembly.

To generate stable cell lines expressing UReKA, PiggyBac transposon plasmids for UReKA expression were constructed by PCR-amplifying the region spanning the start codon of *gvpNJKFGWV* or *gvpA*-IRES-*gvpC* and the end of the hGH polyA from the transient expression plasmids. The amplified regions were then assembled into the PiggyBac transposon backbone (System Biosciences) between a TRE3G promoter (Takara Bio) for doxycycline-inducible expression and a constitutive EF1α core promoter driving either a blasticidin (*BSD)* or hygromycin (*HygR)* resistance genes for selection. A complete list of plasmids used in this study and their sources is provided in Supplementary Table 1. All plasmids were cloned using NEB Stable *E. coli* (New England Biolabs) and verified by Sanger sequencing.

### Expression and Purification of UReKA Gas Vesicles

For in vitro purified protein assays, GVs were collected and purified from confluent *Ana* cultures following previously published protocols^29,39^. Briefly, *Ana* cells were grown in Gorham’s medium supplemented with BG-11 solution (Sigma-Aldrich) and 10 mM sodium bicarbonate at 25 °C under 1% CO_2_ and 100 rpm. shaking, with a 14 h light/10 h dark cycle. Confluent cultures were transferred to sterile separating funnels and left undisturbed for 2–3 days, allowing buoyant *Ana* cells expressing GVs to float and the subnatant was drained. GVs were released by hypertonic lysis using 10% SoluLyse (Amsbio L200125) and 500 mM of sorbitol. Purified GVs were obtained through 3–4 rounds of buoyancy purification at 300g for 4hours per round, where after each round the subnatant was removed and the pellet was resuspended in 6M urea solution buffered with 100 mM of Tris-HCl (pH: 8–8.5) to strip the native outer layer of GvpC. After urea treatment, GVs were either maintained in 6M urea for subsequent GvpC readdition or dialyzed in 4L PBS (Corning) for at least 12 hours at 4 °C for long-term storage. For dialysis, the stripped GVs were loaded into regenerated cellulose dialysis pouches with a 6–8 kDa molecular weight cutoff (Spectrum Labs).

The engineered GvpC variants were expressed and purified following a previously established protocols^29,39^. For protein expression, UReKA GvpC variants were transformed into chemically competent BL21(DE3) cells (Invitrogen) and grown overnight at 37°C for 12–16 hours in 5 ml starter cultures of 2xYT medium supplemented with 50 μg/mL kanamycin. The starter cultures were diluted 1:100 into auto-induction Terrific Broth (Novagen 71491) with 50 μg/mL kanamycin and incubated at 30°C with 250 rpm. shaking for 20-24 hours to induce protein expression. Cells were then harvested by centrifugation at 5,000 rcf and lysed at room temperature using SoluLyse (Amsbio L200125) supplemented with lysozyme (400 μg/mL) and DNase I (10 μg/mL). GvpC inclusion bodies were isolated by centrifugation at 13,000 rcf for 10 minutes, resuspended in solubilization buffer (20 mM of Tris-HCl, 500 mM of NaCl, 6 M of urea, pH 8.0), and incubated with Ni-NTA resin (QIAGEN) for 2 hours at 4°C for His-tag purification. Wash and elution buffers contained the same composition as the solubilization buffer, but with 20 mM and 250 mM imidazole, respectively. The concentration of purified protein was determined using the Bradford Reagent (Bio-rad), and >95% purity was verified by SDS–polyacrylamide gel electrophoresis (PAGE) analysis.

The concentration of Ana GVs was determined by measuring their optical density (OD) at 500 nm (OD_500_) using a Nanodrop spectrophotometer (Thermo Fisher Scientific), with the resuspension buffer as a blank. Based on previous work, a GV concentration of OD_500_ = 1 is ∼184 pM and a gas fraction of 0.0417%. GvpC variants, expressed and isolated from inclusion bodies, were added to stripped *Ana* GVs in 6 M urea in 2-3 fold molar excess after accounting for the 1:15 binding ratio of 3-repeat GvpC:GvpA. The precise quantity of recombinant GvpC (in nanomoles) required for a two-fold stoichiometric excess was calculated using the formula: 2 × OD_500nm_ × 480 nM × volume of GVs (in liters). Recombinant GvpC was refolded onto the surface of the stripped GVs by dialysis in 4L PBS at 4 °C for at least 12 hours. Dialyzed GV samples underwent two or more rounds of buoyancy purification at 300*g* for 3–4 h to remove excess unbound GvpC, where after each round of centrifugation the sample buffer was exchanged into kinase reaction buffer (50mM Tris-HCl, 10mM MgCl_2_, 0.1 mM EDTA, pH 7.5) for further characterization.

### In vitro phosphorylation assay

For all phosphorylation assays, 250 units of PKA (P6000L; NEB; specific activity: 1 pmol/min/unit), 1.5 µg of human CaMKIIα (Sino Biological; specific activity: 160 nmol/min/mg), or 0.1 µg of PKC alpha (Abcam; specific activity: 3,200 nmol/min/mg) per µL of engineered GVs at OD500 = 10 were mixed with 1 mM ATP (Sigma-Aldrich) in NEBuffer™ for Protein Kinases (50 mM Tris-HCl, 10 mM MgCl_2_, 0.1 mM EDTA, 2 mM DTT, 0.01% Brij 35, pH 7.5; NEB) and incubated with slow rotation at 37°C for 10–12 hours, unless otherwise specified. For dephosphorylation assays, 10 units of LPP (P0753L; NEB) per µL of engineered GVs at OD500 = 10 were mixed with PMP Buffer (50 mM HEPES, 100 mM NaCl, 2 mM DTT, 0.01% Brij 35, pH 7.5) and 1 mM MnCl_2_, followed by incubation with slow rotation at 37°C for 10–12 hours.

### Pressurized absorbance spectroscopy

For initial screening of UReKA variants, collapse pressure was determined using previously established protocols^29,39^. Briefly, purified Ana GVs were diluted in experimental buffers to an OD500 of approximately 0.2–0.4, and 400 μL of the diluted sample was loaded into a flow-through quartz cuvette with a 1 cm path length (Hellma Analytics). Hydrostatic pressure was applied using a 1.5 MPa nitrogen gas source, regulated via a single-valve pressure controller (PC series; Alicat Scientific). Optical density (OD500) was continuously measured using a microspectrometer (STS-VIS; Ocean Optics) as pressure increased from 0 to 1 MPa in 50 kPa increments, with a 7-second equilibration period between each measurement. Each dataset was normalized by scaling to the min-max measurement values, and data were fitted using the Boltzmann sigmoid function 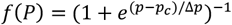 where p_c_, the collapse pressure, represents the midpoint of normalized OD change.The 95% confidence intervals (CIs) were rounded to the nearest integer and are reported in the figures.

### Denaturing polyacrylamide gel electrophoresis (SDS-PAGE and Phos-Tag® Gels)

GV samples incubated with PKA or LPP were concentration-matched at OD500nm = 10 and mixed 1:1 with 2x Laemmli buffer (Bio-Rad), containing SDS and 2-mercaptoethanol. The samples were then boiled at 95°C for 5 minutes and 20 μL of the samples were loaded into either a premade polyacrylamide gel (Bio-Rad) or Phos-Tag® Gel (Fujifilm) immersed in 1x Tris-Glycine-SDS Buffer. In regular SDS-Page gel, 10 uL of Precision Plus ProteinTM Dual Color Standards (Bio-Rad) was loaded as the ladder. Electrophoresis was performed at 120V for 55 minutes, after which the gel was washed in DI water for 15 minutes to remove excess SDS and coomassie-stained for 1 hour on a rocker using the SimplyBlue SafeStain (Invitrogen). The gel was allowed to destain overnight in DI water before imaging the gels using a Bio-Rad ChemiDoc™ imaging system.

### Mass Spectrometry

Proteins were denatured in a buffer containing 8M Urea (U0631, Sigma-Aldrich) and 100 mM Tris pH 8.5. The sample was further reduced by incubation with 10 mM Tris(2-carboxyethyl)phosphine (TCEP, C4706, Sigma) at 37°C for 20 minutes, then alkylated by incubation with 10 mM 2-chloroacetamide (CAA, 79-07-2, Millipore Sigma) in the dark at room temperature for 15 minutes. Subsequently, the sample was digested with 0.1 ug LysC (125-05061, Wako) at 37°C for 4 hours, diluted 4-fold with 100 mM Tris pH 8.5 and CaCl_2_ was added to 1 mM. The sample was then digested with 0.2ug of trypsin (90305, Thermo Fisher Scientific) overnight at 37°C. Samples were acidified by adding trifluoroacetic acid to 0.5%, desalted using Pierce C18 spin columns (89868, Thermo Scientific). After desalting, the peptides were dried and reconstituted in water containing 0.2% formic acid (FA, A11750, Fisher Scientific). and 2% acetonitrile (ACN, A9554, Fisher Scientific) for subsequent LC-MS/MS analysis.

LC-MS/MS analysis was performed with an EASY-nLC 1200 (ThermoFisher Scientific, San Jose, CA) coupled to a Q Exactive HF hybrid quadrupole-Orbitrap mass spectrometer (ThermoFisher Scientific, San Jose, CA). Peptides were separated on an Aurora UHPLC Column (25 cm × 75 μm, 1.6 μm C18, AUR2-25075C18A, Ion Opticks) with a flow rate of 0.35 μL/min for a total duration of 43 min and ionized at 1.7 kV in the positive ion mode. The gradient was composed of 6% solvent B (2 min), 6-25% B (20.5 min), 25-40% B (7.5 min), and 40–98% B (13 min); solvent A: 2% ACN and 0.2% FA in water; solvent B: 80% ACN and 0.2% FA. MS1 scans were acquired at the resolution of 60,000 from 375 to 1500 m/z, AGC target 3e6, and maximum injection time 15 ms. The 12 most abundant ions in MS1 scans are selected for fragmentation via higher-energy collisional dissociation (HCD) with a normalized collision energy (NCE) of 28. MS2 scans were acquired at a resolution of 30,000, AGC target 2e5 and a maximum injection time 120 ms. Dynamic exclusion was set to 30 s and ions with charge +1, +7, +8 and >+8 were excluded. The temperature of ion transfer tube was 275°C and the S-lens RF level was set to 55.

RAW files were searched with Proteome Discoverer SEQUEST (version 2.5, Thermo Scientific) against the fasta file containing *Anabaena flos-aquae* GVPC, GVPAand mutant GVPC protein sequence. Trypsin/P was set as the digestion enzyme, allowing a maximum of two missed cleavages. Dynamic modifications were set to oxidation on methionine (M, +15.995 Da), deamidation on asparagine and glutamine (N and Q, +0.984 Da), phosphorylation on serine, threonine and tyrosine (S, T and Y, +79.966 Da) and protein N-terminal acetylation (+42.011 Da) and Met-loss (-131.040 Da). Carbamidomethylation on cysteine residues (C, +57.021 Da) was set as a fixed modification. The maximum parental mass error was set to 10 ppm, and the MS2 mass tolerance was set to 0.05 Da. The relative abundance of parental peptides was calculated by integration of the area under the curve of the MS1 peaks using the Minora LFQ node. The maximum false peptide discovery rate was specified as 0.01 using the Percolator Node validated by q-value. The IMP-ptmRS node within the Proteome Discoverer was used to calculate PTM site probabilities. The RAW data have been deposited to the ProteomeXchange Consortium via the PRIDE partner repository with the dataset identifier PXD071157.^61^ Spectral annotation was generated by the Interactive Peptide Spectral Annotator (IPSA).^62^

### HEK293T cell culture and transient transfection

HEK293T cells (American Type Culture Collection (ATCC), CLR-2316) were seeded in 12-well plates at a density of 1.5 × 10^5^ cells per well and maintained at 37 °C with 5% CO2 in a humidified incubator. Cells were cultured in 1 mL of DMEM (Corning, 10-013-CV) with 10% FBS (Gibco), 10 mM HEPES (Cytia) and 1× penicillin–streptomycin 24 hours before transfection. Transient transfection mixtures were created by mixing 2 µg of plasmid mixture with Transporter− 5 (Polysciences 26008) at 1:4 DNA/Transporter ratio. The mixture was incubated for 20 minutes at room temperature and added drop-wise to HEK293T cells. Media was changed after 12–16 hours and daily thereafter. For the transient expression of UReKA and dUReKA, the mutant control, 280 fmol of the *gvpA*-IRES-*gvpC* plasmid and 70 fmol of the plasmid encoding *gvpNJKFGWV* was added into the DNA mixture, and pUC19 plasmid DNA was supplemented to make the total amount of DNA 2 µg. For PKA activation study, either 35 fmol of the *PRKACα* were added to the DNA mixture to overexpress the catalytic subunit or the media were supplemented with 20uM forskolin (MedChemExpress) and 500uM of 3-isobutyl-1-methylxantine (Sigma-Aldrich) 12-hr prior harvesting the cells to saturate the PKA activity level. For PKA inhibition study, 50uM of H-89 dihydrocloride (Tocris) was supplemented to the media for at least 3hrs. Transfected cells were assayed 72 hours after the transfection.

### Purified GV and in vitro Ultrasound Imaging

After 3 days of expression, cells were dissociated using Trypsin/EDTA (Corning 25-053-CI), centrifuged at 300 rcf for 6 minutes at room temperature and resuspended in PBS. The cells were counted using an automated cell counter (Countess™ 3, Thermo Fisher) and all the samples were concentration-matched to 10 million cells per milliliter in PBS. Imaging phantoms were prepared with 1% agarose (w/v, Lonza, #50070) in PBS. For ultrasound imaging, either purified GVs or cells were diluted 1:1 with the 1% agarose to result in a final concentration of OD1 GVs or 5 million cells per milliliter, respectively, before loading into their respective phantoms or injected into an acoustically transparent tubing for continuous kinetics experiments. All the samples were placed in the same buffer as the phantom during the imaging session. For kinetic measurements, the samples were maintained at 37°C by a custom water bath.

Imaging was performed using a Verasonics Vantage programmable ultrasound scanning system and a L22-14vX 128-element linear array Verasonics transducer, with a specified pitch of 0.1 mm, an elevation focus of 8 mm, an elevation aperture of 1.5mm and a center frequency of 18.5 MHz with 67% -6 dB bandwidth. For nonlinear image acquisition, a custom cross-amplitude modulation (xAM) sequence detailed in an earlier study^31^ was used with an xAM angle (θ) of 19.5°, an aperture of 65 elements, and a transmitting frequency at 15.625 MHz. The center of the sample wells was placed at a depth of 5 mm with a conventional ray-line scanning B-mode pulse sequence with parabolic focusing at 10 mm and an aperture of 40 elements. The focus was set to be far from the sample position to reduce the acoustic pressure to avoid collapsing the samples. The transmitted pressure at the sample position at 5 mm was calibrated using a Precision Acoustics fiber-optic hydrophone system. Each image was an average of 50 accumulations.

B-mode images were acquired at a transmit voltage of 1.6V (50 kPa), and an automated voltage ramp imaging script (programmed in MATLAB) was used to conduct xAM acquisitions at different acoustic pressures. For all samples for characterization, an xAM voltage ramp sequence from 3V (219 kPa) to 8V (939 kPa) in 0.5V step increments was used. Samples were subjected to complete collapse at 25V with the B-mode sequence for 10 seconds, and the subsequent post-collapse xAM images acquired at the same voltage steps were used as the blank value for data processing. For purified GV samples, B-mode images at the beginning of the pressure ramp and at the end of the post-collapse ramp were also acquired for concentration normalization. For continuous kinetics monitoring of dephosphorylation in purified GVs, xAM image was taken at 5V (489 kPa) at 0.5 frames per minute for 150 minutes.

## ACKNOWLEDGEMENTS

The authors thank R. Nayak for providing calibration data for the ultrasound transducer used for imaging. Mass spectrometry was done in the Protein Exploration Laboratory at Caltech. This research was supported by the Chan-Zuckerberg Initiative and the National Institutes of Health (R01EB018975 to M.G.S.). J.W.Y. was supported by the Biotechnology Leadership pre-doctoral Training Program Fellowship (Rosen Bioengineering Center). M.G.S. is an investigator of the Howard Hughes Medical Institute.

## AUTHOR CONTRIBUTIONS

J.W.Y., Z.J., and M.G.S. conceived and designed the study. J.W.Y., and Z.J. designed and planned experiments. J.W.Y., and T.W. conducted the experiments. J.W.Y., and Z.J. wrote the scripts for ultrasound imaging and data processing. J.W.Y. analyzed the data. J.W.Y. wrote the manuscript, with input from all authors. M.G.S. supervised the research. All authors have given approval to the final version of the manuscript.

