## Supplementary Figures for "Ultrasonic Reporter of Kinase Activity"

### Supplementary Materials

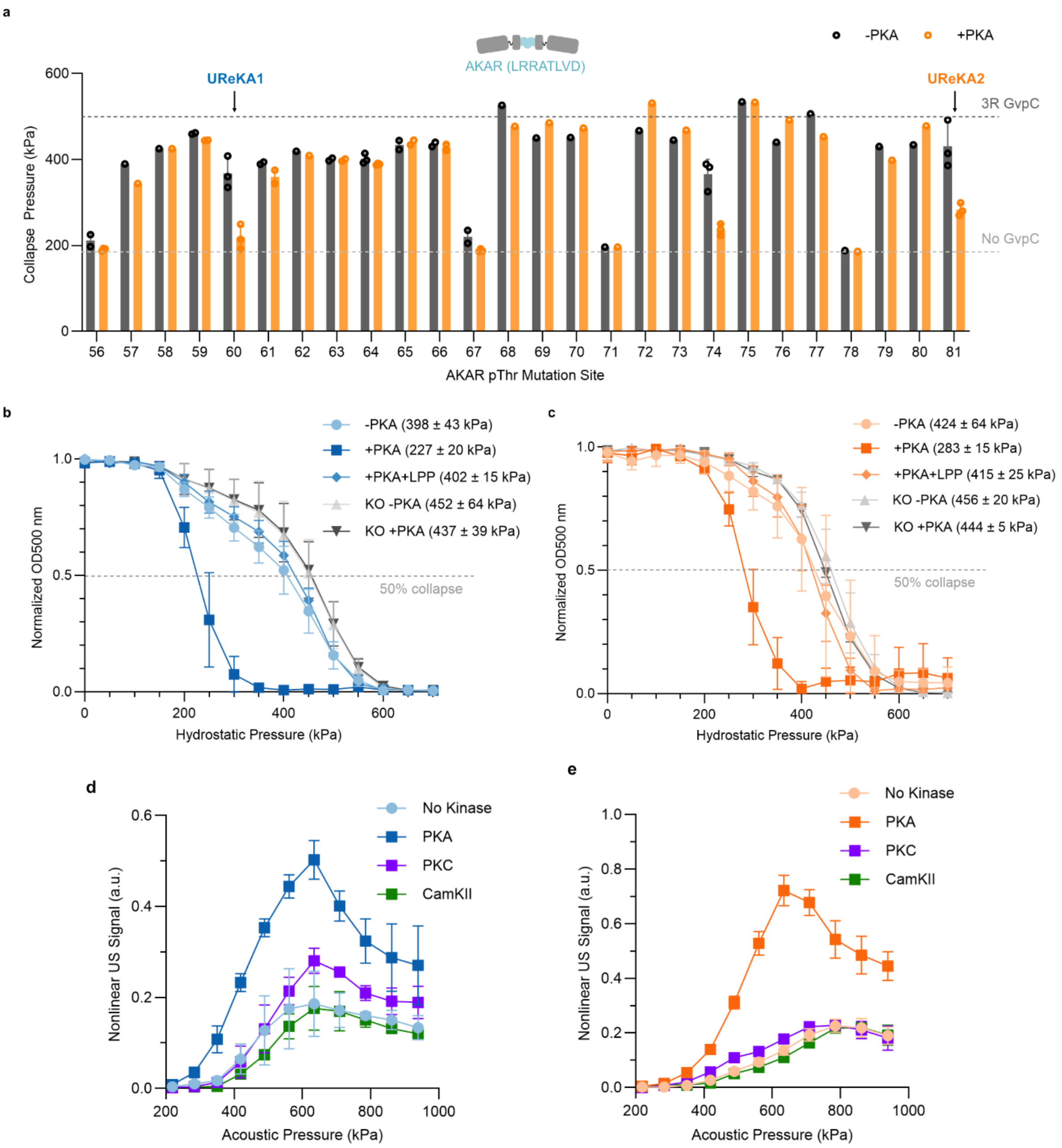

**Supplementary Figure 1: Other Characterization of UReKA.** a) Hydrostatic collapse pressure screening of UReKA variants with AKAR sequence at various phosphothreonine (pThr) position in GvpC. (n=3 for 60, 74, 81, n=1 or 2 for others). Midpoint collapse pressure for each variant was determined from fitting a Boltzmann sigmoid function. Samples at $OD_{500}$ 10 were incubated with 1mM ATP and with (orange) or without (black) 5,000 units of cAMP-activated PKA at 37 ° C. Error bar indicates SEM based on curve fitting. Dashed lines indicate the mean signal of GVs with 3R GvpC (black) or without GvpC (gray). b) Normalized $OD_{500}$ of UReKA1 at various conditions as a function of hydrostatic pressure without PKA, with PKA, with PKA and LPP, and T/A knockout (KO) used as a control. (n= 3 biological replicates) c) Normalized $OD_{500}$ of UReKA2 at various conditions as a function of hydrostatic pressure without PKA, with PKA, with PKA and LPP, and T/A knockout (KO) used as a control. (n= 3 biological replicates) d) Nonlinear ultrasound signal normalized by B-mode intensity as a function of acoustic pressure for UReKA1, incubated with PKA (blue), PKC (purple), and CamKII (green). (n=3) e) Nonlinear ultrasound signal normalized by B-mode intensity as a function of acoustic pressure for UReKA2, incubated with PKA (blue), PKC (purple), and CamKII (green). (n=3)

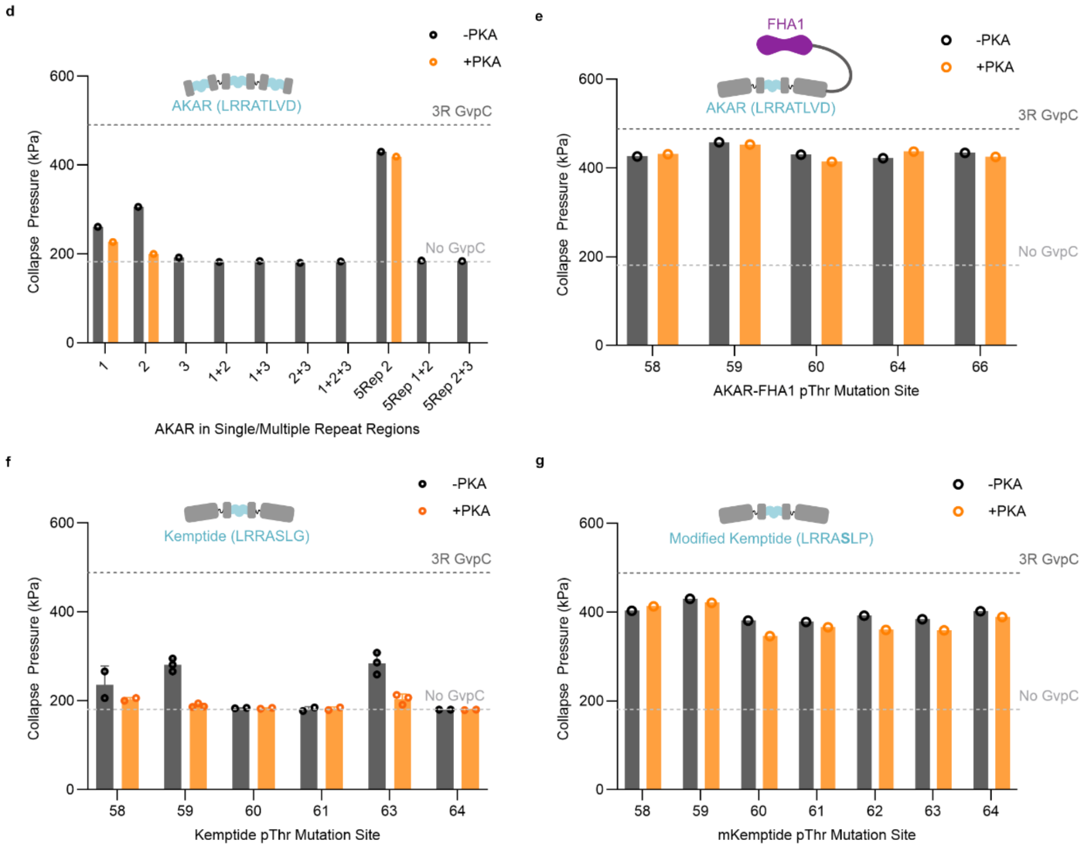


**Supplementary Figure 2: Other UReKA Variant Screenings.** a)-d) Midpoint collapse pressure for each variant was determined from fitting a Boltzmann sigmoid function. Samples at $OD_{500}$ 10 were incubated with 1 mM ATP and with (orange) or without (black) 5,000 units of cAMP-activated PKA at 37 °C. Error bar indicates SEM based on curve fitting. Dashed lines indicate the mean signal of GVs with 3R GvpC (black) or without GvpC (gray). a) Hydrostatic collapse pressure screening of AKAR sequence in various repeats of GvpC (1=27, 2=60, 3=93, n=1) b) Hydrostatic collapse pressure screening of AKAR-FHA1 UReKA variants (n=1) c) Hydrostatic collapse pressure screening of Kemptide-UReKA variants (n=3 for 59 and 63, n=2 for others) d) Hydrostatic collapse pressure screening of modified Kemptide-UReKA variants (n=1)

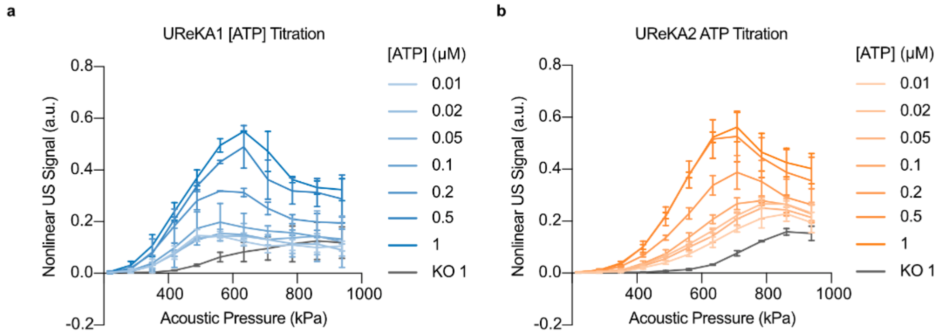


**Supplementary Figure 3: ATP titration xAM response.** a) Nonlinear ultrasound signal of UReKA1 (blue) incubated with PKA and various concentrations of ATP from 10nM to 1mM with respective T/A KO control (gray). b) Nonlinear ultrasound signal of UReKA2 (orange) incubated with PKA and various concentrations of ATP from 10nM to 1mM with respective T/A KO control (gray).

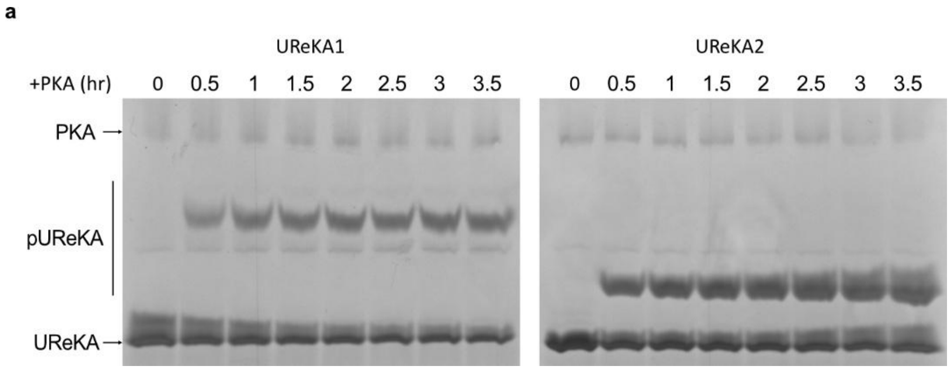


**Supplementary Figure 4: Pseudo-quantitative phosphorylation kinetics analysis.** a) Coomassie Brilliant Blue-stained SuperSep™ Phos-Tag® SDS-Page gel image of UReKA (bottom band) at various incubation timepoint with PKA (top band). Band intensity between the top and bottom bands indicate phosphorylated fraction of UReKA GvpC.

a

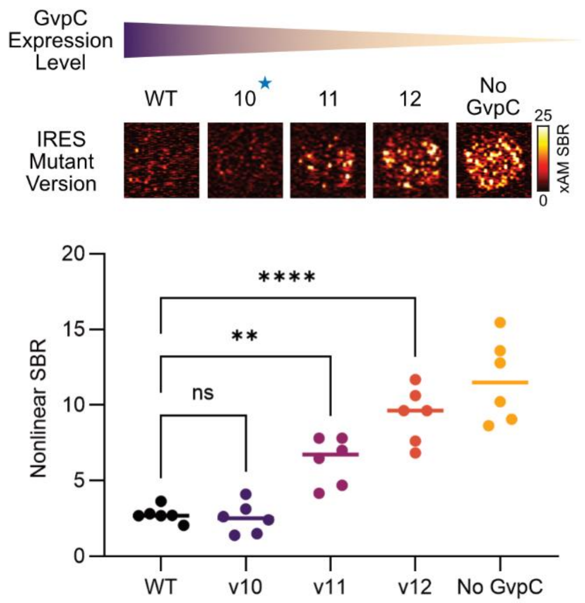


**Supplementary Figure 5: Supplementary Figure 5: UReKA variant screening with mutated IRES.** Representative nonlinear xAM image at 419 kPa (top) of various UReKA GVs expressed with mutant IRES from v9-12 upstream of GvpC with respective SBR (below). n=6.
